# A mutagenesis screen for essential plastid biogenesis genes in human malaria parasites

**DOI:** 10.1101/401570

**Authors:** Yong Tang, Thomas R. Meister, Marta Walczak, Michael J. Pulkoski-Gross, Sanjay B. Hari, Robert T. Sauer, Katherine Amberg-Johnson, Ellen Yeh

## Abstract

Endosymbiosis has driven major molecular and cellular innovations. *Plasmodium* spp. parasites that cause malaria contain an essential, non-photosynthetic plastid, the apicoplast, which originated from a secondary (eukaryote-eukaryote) endosymbiosis. To discover organellar pathways with evolutionary and biomedical significance, we performed a mutagenesis screen for essential genes required for apicoplast biogenesis in *P. falciparum*. *Apicoplast-minus* mutants were isolated using a chemical rescue that permits conditional disruption of the apicoplast and a new fluorescent reporter for organelle loss. Five candidate genes were validated (out of 12 identified), including a TIM-barrel protein that likely derived from a core metabolic enzyme but evolved a new activity. Our results demonstrate the first forward genetic screen to assign essential cellular functions to unannotated *P. falciparum* genes. A putative TIM-barrel enzyme and other newly-identified apicoplast biogenesis proteins open opportunities to discover new mechanisms of organelle biogenesis, molecular evolution underlying eukaryotic diversity, and drug targets against multiple parasitic diseases.

## Introduction

*Plasmodium* spp. which cause malaria and related apicomplexan parasites are important human and veterinary pathogens. These disease-causing protozoa are highly divergent from well-studied model organisms that are the textbook examples of eukaryotic biology, such that parasite biology often reveals striking eukaryotic innovations. The apicoplast, a nonphotosynthetic plastid found in apicomplexa, is one such “invention” (Kohler et al., 1997; McFadden et al., 1996). These intracellular parasites evolved from photosynthetic algae that acquired their plastids through secondary endosymbiosis (van Dooren and Striepen, 2013). During secondary endosymbiosis, a chloroplast-containing alga, itself the product of primary endosymbiosis, was taken up by another eukaryote to form a secondary plastid (Gibbs, 1978). Despite the loss of photosynthesis in apicomplexans, the apicoplast contains several prokaryotic metabolic pathways, is essential for parasite replication during human infection, and is a target of antiparasitic drugs (Dahl and Rosenthal, 2007; Fichera and Roos, 1997; Ralph et al., 2004; Sheiner et al., 2013).

There are many traces of the apicoplast’s quirky evolution in its present-day cell biology, in particular the pathways for its biogenesis. Like other endosymbiotic organelles, the single apicoplast cannot be formed *de novo* and must be inherited by its growth, division, and segregation into daughter cells. The few molecular details we have about apicoplast biogenesis hint at the major innovations that have occurred in the process of adopting and retaining this secondary plastid. The apicoplast is bound by four membranes acquired through successive endosymbioses, such that apicoplast proteins transit through the ER and use a <u>s</u>ymbiont-derived <u>E</u>RAD-like <u>ma</u>chinery (SELMA) to cross the two new outer membranes (Agrawal et al., 2009). Curiously, autophagy-related protein 8 (Atg8), a highly-conserved eukaryotic protein and key marker of autophagosomes, localizes to the apicoplast and is required for its inheritance in *Plasmodium* and related apicomplexan parasite *Toxoplasma gondii* (Kitamura et al., 2012; Leveque et al., 2015; Walczak et al., 2018). While SELMA and *Pf*Atg8 are clear examples of molecular evolution in action, other novel or repurposed proteins required for apicoplast biogenesis remain undiscovered.

So far, new apicoplast biogenesis proteins have primarily been discovered through candidate approaches. SELMA was first identified as homologs of the ER-associated degradation (ERAD) machinery encoded in the nucleomorph, the remnant nucleus of the eukaryotic symbiont found in some algal secondary plastids (Sommer et al., 2007). Nuclear-encoded versions of SELMA containing apicoplast targeting sequences were then detected in the genomes of apicomplexan parasites (which lack a nucleomorph) (Spork et al., 2009). Since the apicoplast proteome is enriched in biogenesis functions, several apicoplast-targeted proteins of unknown function have also been shown to be required for its biogenesis (Sheiner et al., 2011) (unpublished data). ATG8’s novel apicoplast function was discovered serendipitously by its unexpected localization on the apicoplast instead of autophagosomes (Kitamura et al., 2012). Though candidate approaches have yielded new molecular insight (Biddau et al., 2018; Fellows et al., 2017; Sheiner et al., 2015), in general they are indirect and may bias against novel pathways.

In blood-stage *P. falciparum*, a remarkable method to chemically rescue parasites that have lost the apicoplast has paved the way for functional screens (Gisselberg et al., 2018; Wu et al., 2015; Yeh and DeRisi, 2011). Incredibly, addition of isopentenyl diphosphate (IPP) to the growth media is sufficient to reverse growth inhibition caused by apicoplast loss, since it is the only essential metabolic product of the apicoplast in the blood stage. Taking advantage of the apicoplast chemical rescue, we recently took the first unbiased approach to discover a new apicoplast biogenesis protein (Amberg-Johnson et al., 2017). We first screened for small molecule inhibitors that specifically disrupt apicoplast biogenesis in *P. falciparum*. Subsequent target identification led us to a membrane metalloprotease, *Pf*FtsH1, with an unexpected role in apicoplast biogenesis. This chemical genetic screen has the advantage of unbiased sampling of druggable targets in apicoplast biogenesis pathways. Unfortunately, it lacks throughput given the painstaking process of mapping inhibitors to their molecular targets.

Forward genetic screens are powerful approaches to uncover novel cellular pathways, such as those required for apicoplast biogenesis. Recently, genome-scale deletion screens performed in several apicomplexan parasites have uncovered a plethora of essential genes of unknown function (Bushell et al., 2017; Sidik et al., 2016; Zhang et al., 2018). Several challenges impede large-scale functional analysis of these essential genes. First, targeted gene modifications are still slow and labor-intensive in *P. falciparum,* the deadliest of the human malaria species and the only *Plasmodium* spp. that can be cultured *in vitro* (Schuster, 2002; Trager and Jensen, 1976). Second, efficient methods for generating conditional mutants, such as RNAi or CRISPRi systems, are lacking in all apicomplexan organisms (de Koning-Ward et al., 2015). Finally, high-throughput, single-cell phenotyping for important functions need to be developed (Matz and Kooij, 2015). Overcoming these significant limitations, we designed a forward genetic screen using chemical mutagenesis, apicoplast chemical rescue, and a fluorescent reporter for apicoplast loss to identify essential apicoplast biogenesis genes in blood-stage *P. falciparum*. The screen identified known and novel genes required for apicoplast biogenesis and is the first forward genetic screen for essential pathways in *P. falciparum*.

## Results

### A conditional fluorescent reporter enables single-cell phenotyping of apicoplast loss in *P. falciparum*

To isolate rare *P. falciparum* mutants that have lost their apicoplast due to biogenesis defects, we set out to design a live-cell reporter for apicoplast loss. The apicoplast contains a prokaryotic caseinolytic protease (Clp) system composed of a chaperone ClpC that recognizes and unfolds substrates and a protease ClpP that degrades the recognized substrates (El Bakkouri et al., 2010; Florentin et al., 2017; Rathore et al., 2010). We hypothesized that (1) in the presence of a functional apicoplast, ClpCP could be co-opted to degrade and turn ‘off’ a fluorescent reporter, whereas (2) loss of the apicoplast would result in loss of ClpCP activity and therefore turn ‘on’ the reporter (Figure 1A).

**Figure 1.**
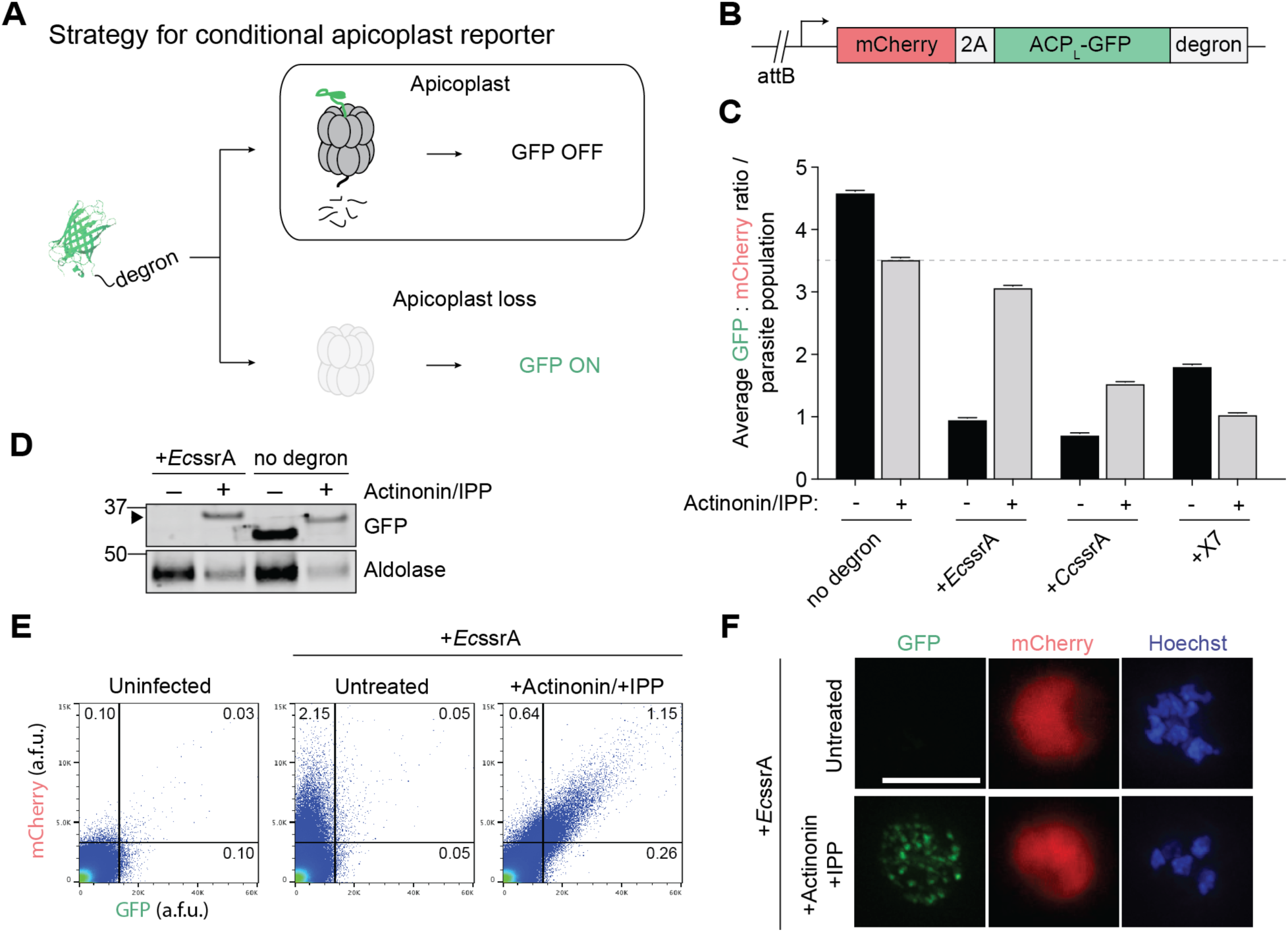
A fluorescent reporter for apicoplast loss in *P. falciparum* is specifically degraded in the apicoplast. (A) Strategy for a conditional GFP reporter that fluoresces upon apicoplast loss. (B) Reporter construct for expression of apicoplast-targeted GFP (ACP_L_-GFP) and cytoplasmic mCherry used for integration into *P. falciparum* Dd2^attB^ parasites. (C) Ratio of GFP: mCherry fluorescence in untreated versus actinonin/IPP-treated parasites expressing ACP_L_-GFP-degron. Data are shown as mean ± s.e.m. (n=2). See also Figure S1. (D) GFP protein levels in untreated versus actinonin/IPP treated parasites expressing GFP-*Ec*ssrA. Higher molecular weight species in GFP blot from unprocessed ACP leader peptide (ACP_L_) is indicated with black arrowhead. (E) Flow cytometry plots showing mCherry and GFP fluorescence in untreated versus actinonin/IPP treated parasites expressing GFP-*Ec*ssrA. Uninfected red blood cells (RBCs) are gated away (lower left quadrants) and the % mCherry+, GFP+ parasites in the gated populations relative to the total number of cells quantified are indicated. (F) Representative live-cell fluorescent images of untreated and actinonin/IPP treated parasites expressing mCherry and GFP-*Ec*ssrA. Hoechst stains for parasite nuclei. Scale bar 5 µm.

Clp substrates are typically recognized by unstructured degron sequences, the best studied of which is a ssrA peptide appended to the C-terminus of translationally-stalled proteins (Gottesman et al., 1998; Keiler et al., 1996). However, the substrate specificity of the apicoplast ClpCP system has yet to be defined. Reasoning that *Pf*ClpCP homolog might recognize similar degrons as bacterial or algal Clp systems, we tested two degrons recognized by *E. coli* ClpXP, *Ec*ssrA and X7, and the predicted ssrA peptide in the red alga *Cyanidium caldarium, Cc*ssrA (Table S1) (Flynn et al., 2001; Hoskins and Wickner, 2006; Hudson and Williams, 2015). To assess their functionality in *P. falciparum*, the *C*-terminus of an apicoplast-targeted GFP (ACP_L_-GFP) was modified with each of the degrons (Figure 1B). A cytosolic mCherry marker was also expressed on the same promoter as ACP_L_-GFP via a T2A ‘skip’ peptide to normalize apicoplast GFP levels to stage-specific promoter expression (Wagner et al., 2013). Each construct was then integrated into an ectopic *att*B locus in Dd2^attB^ parasites to generate the reporter strains (Nkrumah et al., 2006).

For each reporter strain, the ratio of GFP:mCherry fluorescence (as detected by flow cytometry) was assessed in untreated parasites containing an intact apicoplast, designated *apicoplast(+)*, and in parasites treated with actinonin (which causes apicoplast loss via inhibition of *Pf*FtsH1) and rescued with IPP rendering them *apicoplast(-)*. In the absence of a degron, the GFP:mCherry ratio decreased modestly in *apicoplast(-)* parasites compared to *apicoplast(+)* (Figure 1C and S1A). Though GFP was present in both populations, live fluorescence microscopy confirmed its localization to a branched structure characteristic of the apicoplast in *apicoplast(+)* parasites and to dispersed punctae as previously reported in *apicoplast(-)* parasites (Figure S1B) (Yeh and DeRisi, 2011). Addition of each of the 3 degrons to the C-terminus of ACP_L_-GFP caused a 60-84% decrease in the GFP:mCherry ratio in *apicoplast(+)* parasites, consistent with specific degradation of GFP (Figure 1C). In ACP_L_-GFP-*Ec*ssrA and ACP_L_-GFP-*Cc*ssrA populations, the GFP:mCherry ratio recovered to 88% and 44% relative to the ACP_L_-GFP population in *apicoplast(-)* parasites, respectively (Figure 1C, 1E, and S1C). Unexpectedly, ACP_L_-GFP-X7 populations showed further reduction of GFP:mCherry in *apicoplast(-)* parasites (Figure 1C and S1C). Notably, cytoplasmic mCherry levels were similar between *apicoplast(+)* and *apicoplast(-)* parasites in all reporter strains, suggesting that the differences in GFP levels were not due to altered expression levels (Figures S1D). Of the 3 degrons tested, both the *Ec*ssrA and *Cc*ssrA peptide caused apicoplast-specific GFP degradation as designed.

We further characterized ACP_L_-GFP-*Ec*ssrA, since the greatest recovery of GFP fluorescence following apicoplast loss was observed with this reporter (Figure 1C). Consistent with degradation of GFP in the apicoplast dependent on *Ec*ssrA peptide, ACP_L_-GFP-*Ec*ssrA protein was only detected in *apicoplast(-)* parasites while unmodified ACP_L_-GFP was detected in both *apicoplast(+)* and *(-)* parasites (Figure 1D). Of note, cleavage of the apicoplast-targeting ACP_L_ sequence does not occur in *apicoplast(-)* parasites, resulting in a ACP_L_-GFP protein of higher molecular weight compared to *apicoplast(+)* parasites (Yeh and DeRisi, 2011). Flow cytometry and live fluorescence microscopy confirmed that *apicoplast(+)* parasites displayed only cytosolic mCherry fluorescence, while *apicoplast(-)* parasites displayed both cytosolic mCherry and dispersed, punctate GFP fluorescence (Figure 1E and 1F). Taken together, these results demonstrate that ACP_L_-GFP-*Ec*ssrA serves as a specific reporter for apicoplast loss in *P. falciparum.*

### A phenotypic screen isolates a collection of apicoplast(-) mutant clones

Next we used the ACP_L_-GFP-*Ec*ssrA reporter strain to perform a phenotypic screen for *apicoplast(-)* mutants (Figure 2A). Ring-stage parasites were mutagenized with the alkylating agents ethyl methanesulfonate (EMS) or *N*-ethyl-*N*-nitrosourea (ENU) to generate a diverse population of parasites, some of which harbor mutations in apicoplast biogenesis genes rendering them *apicoplast(-)*. To rescue the lethality of apicoplast loss, mutagenized parasites were supplemented with 200 µM IPP in the growth media. A control group of non-mutagenized parasites was also cultured with IPP to assess whether *apicoplast(-)* mutants naturally occurred over time in the absence of selective pressure to maintain the organelle.

**Figure 2.**
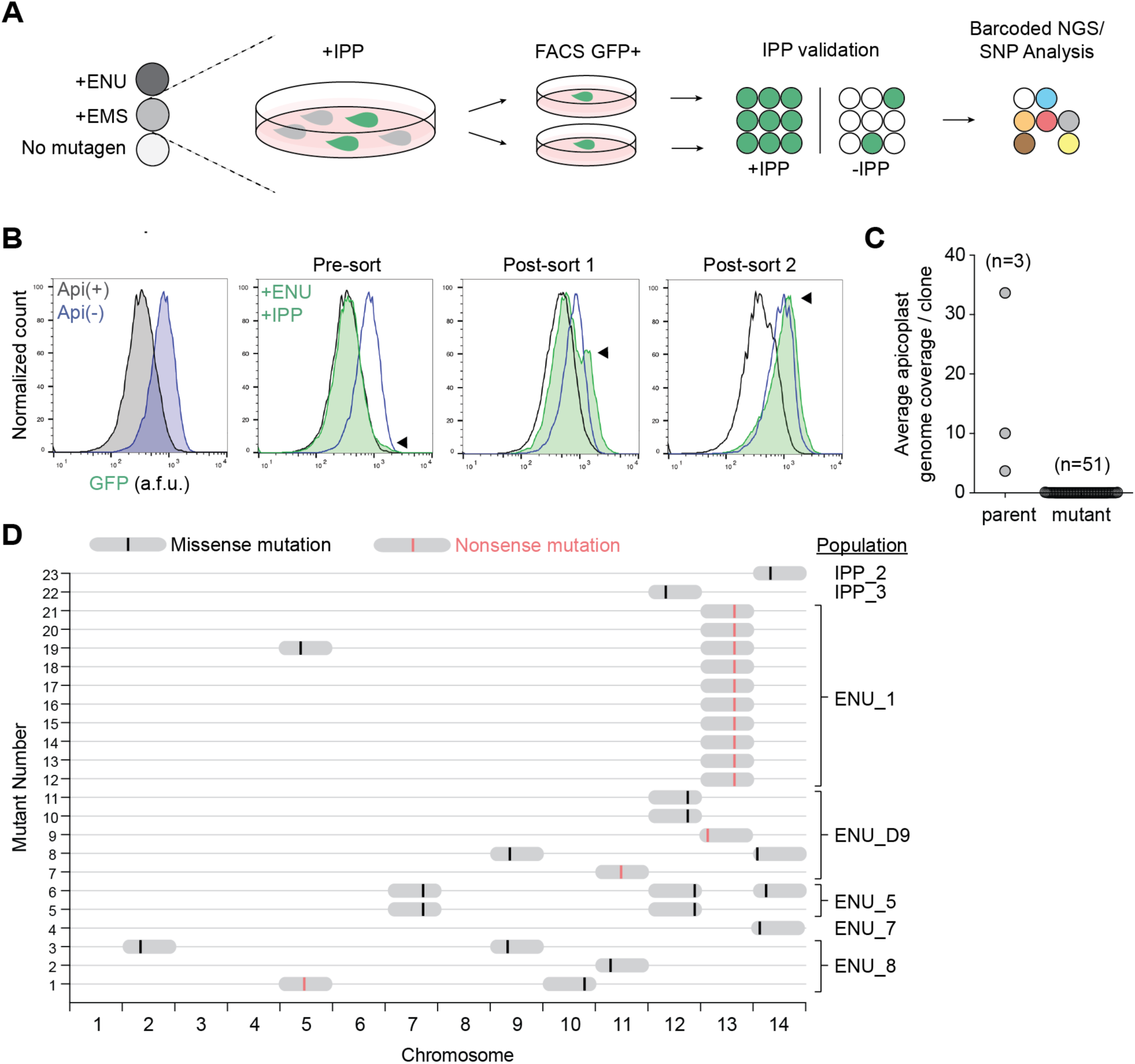
A screen for *apicoplast(-)* mutant clones identifies candidate biogenesis genes. (A) Schematic for selection and characterization of *apicoplast(-)* mutant clones. (B) Representative flow cytometry histograms of ENU_1 population (shown in D) GFP fluorescence over the course of successive fluorescence-activated cell sorts. Distribution of GFP fluorescence for pure populations of *apicoplast(+)* and *apicoplast(-)* parasites are shown for comparison in grey and blue, respectively. (C) Apicoplast genome coverage in parental strains and mutant clones from whole-genome sequencing. (D) Genomic loci of SNVs identified in coding regions of mutant clones. The mutagenized populations from which each clone was derived is indicated. For specific mutations, see also Table 1 and Table S2.

After two replication cycles to allow for initial apicoplast loss, mCherry-expressing parasites displaying GFP fluorescence > ∼70% percentile were isolated by fluorescence-activated cell sorting (FACS). Selected mCherry(+) and GFP(+) parasites were then allowed to propagate to a detectable parasitemia before being subjected to a another round of FACS. In one exemplary ENU-mutagenized population, a distinct population of GFP-positive parasites began to emerge after just two rounds of FACS (Figure 2B). Other populations showed enrichment beginning after three rounds of FACS. No significant enrichment of GFP signal was observed when parasites were (1) grown with IPP over time without FACS or (2) grown without IPP and subjected to FACS, suggesting that we specifically enriched for *apicoplast(-)* mutants. Parasites from the final enriched *apicoplast(-)* populations were individually sorted to generate *apicoplast(-)* clones derived from single parasites. Each clonal population was then re-checked for IPP-dependent growth and GFP fluorescence.

### Whole-genome sequencing identifies 12 candidate genes required for apicoplast biogenesis

To determine the genetic basis of apicoplast loss, we sequenced the genomes of 51 *apicoplast(-)* clones (mutagenized = 40, non-mutagenized = 11) and three parent populations. Sequenced reads were mapped to the genome of *Plasmodium falciparum* 3D7 (version 35) with an average read depth of 26 for all samples. Notably, while the apicoplast genome was sequenced at an average depth of 16 reads in the parent populations, it was only detected with an average of 0.02 reads in *apicoplast(-)* clones (Figure 2C). Since the organellar genome is a marker for the apicoplast, the absence of the apicoplast genome confirmed the loss of the apicoplast in the sequenced clones (Amberg-Johnson et al., 2017; Yeh and DeRisi, 2011).

We next performed single nucleotide variant (SNV) analysis. A raw variant list was generated for each sample by comparison to the reference 3D7 genome and included SNVs found in parent populations at ≥5% allele frequency (minimum one read) and in *apicoplast(-)* clones at ≥90% (minimum 5 reads). Any SNV identified in *apicoplast(-)* clones that was also identified in any of the three parent populations was removed. We also filtered out SNVs detected in non-coding regions or resulting in synonymous amino acid changes in coding regions. Finally, SNVs identified in hypervariable regions of the genome (including the rifin, stevor and EMP gene families) and/or previously annotated in the PlasmoDB single nucleotide polymorphism (SNP) database were excluded. After these filtering steps, 23 *apicoplast(-)* clones had at least one but no more than three SNVs that differed from the parent populations (Figure 2D; Table S2).

Since genes required for apicoplast biogenesis ought to be essential, we used essentiality data from literature or whole-genome deletion screens performed in blood-stage *P. falciparum* and *P. berghei* to prioritize gene candidates (Bushell et al., 2017; Zhang et al., 2018). Of 18 unique SNVs identified, 12 were in genes categorized as “essential” in blood-stage *P. falciparum* and/or *P. berghei* (Table 1 and Table S2). Although *Pf*FtsH1 (Pf3D7_1239700) is categorized as “dispensable” in the *P. falciparum* deletion screen, it has been shown experimentally to be essential (Zhang et al., 2018). Overall, a mutation in one of these 12 essential gene candidates was identified in each of the 23 *apicoplast(-)* clones, consistent with the mutation causing apicoplast loss. Potentially disruptive mutations included a I437S variant in the known apicoplast biogenesis protein (*Pf*FtsH1), truncation of autophagy-related protein 7 (*Pf*Atg7) likely required for a cytoplasmic pathway for apicoplast biogenesis, and truncations of three proteins of unknown function (Pf3D7_0518100, Pf3D7_1305100, Pf3D7_1363700) (Table 1). The remaining candidates contained point mutations and had no known prior function in apicoplast biogenesis or localization to the organelle.

**Table 1.** List of top candidate genes from screen. Candidates were rank ordered based on type of mutation and previous literature on candidate genes. Note that the first 2 digits of the Pf gene ID indicate the chromosome on which the gene is located.

| Gene ID | Annotation | Variant | Essential in *P. falciparum* piggyBac screen<sup>a</sup> | Essential in *P. berghei* screen<sup>b</sup> | Essential in other literature |
| --- | --- | --- | --- | --- | --- |
| **Pf3D7_1239700** | ATP-dependent zinc metalloprotease FtsH1 | Ile437Ser | No | ND | Yes<sup>c</sup> |
| **Pf3D7_1126100** | autophagy-related protein 7, putative | Gln719* | Yes | ND | Yes<sup>d</sup> |
| **Pf3D7_0518100** | RAP protein, putative | Leu141* | No | Yes | ND |
| **Pf3D7_1305100** | conserved Plasmodium protein, unknown function | Lys85* | Yes | ND | ND |
| **Pf3D7_1363700** | conserved Plasmodium protein, unknown function | Tyr466* | Yes | Yes | ND |
| **Pf3D7_0913500** | protease, putative | Ser347Gly | Yes | ND | ND |
| **Pf3D7_1408700** | conserved Plasmodium protein, unknown function | Ile4809Val | No | Yes | ND |
| **Pf3D7_1114900** | conserved Plasmodium protein, unknown function | Lys932Ile | Yes | Yes | ND |
| **Pf3D7_0910100** | exportin-7, putative | Ser102Pro | Yes | Yes | ND |
| **Pf3D7_1417800** | DNA replication licensing factor MCM2 | Thre584Ile | Yes | Yes | ND |
| **Pf3D7_1433800** | conserved Plasmodium protein, unknown function | Asp431Val | Yes | ND | ND |
| **Pf3D7_1203300** | conserved Plasmodium protein, unknown<br>function | Lys473Thr | Yes | ND | ND |

<sup>a</sup> From *P. falciparum* piggyBac screen dataset (Zhang et al., 2018)
<sup>b</sup> From *P. berghei* deletion screen dataset (Bushell et al., 2017)
<sup>c</sup> (Amberg-Johnson et al., 2017)
<sup>d</sup> (Walker et al., 2013)

### The Ile437Ser variant causes loss of *Pf*FtsH1 activity *in vitro*

*Pf*FtsH1 is an apicoplast membrane metalloprotease that was previously identified in a chemical genetic screen as the target of actinonin, an inhibitor that disrupts apicoplast biogenesis. Subsequent knockdown of *Pf*FtsH1demonstrated that it is required for apicoplast biogenesis (Amberg-Johnson et al., 2017). We hypothesized that the I437S variant identified in our screen disrupted *Pf*FtsH1 function, leading to apicoplast loss. *Pf*FtsH1 contains both an ATPase and peptidase domain. To test the effect of I437S on enzyme activity, we compared the activity of I437S with that of wild-type (WT) enzyme, an ATPase-inactive E249Q variant, and a peptidase-inactive D493A variant (Figure 3A). All enzymes were purified without the transmembrane domain as previously described (Amberg-Johnson et al., 2017).

**Figure 3.**
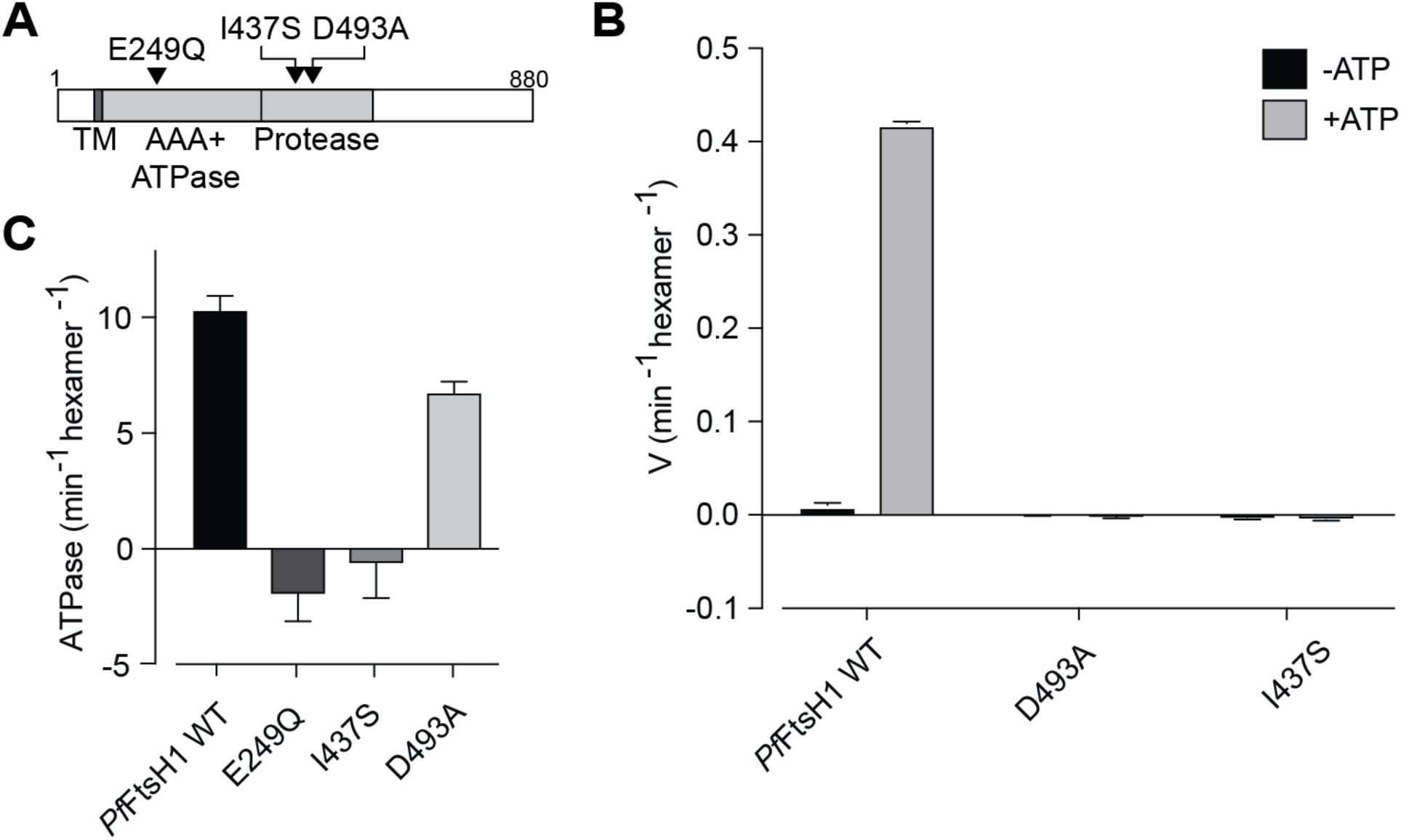
The I437S variant disrupts *Pf*FtsH1 protease and ATPase activity *in vitro*. (A) Domain organization of *Pf*FtsH1 showing location of I437S. The known ATPase-inactive (E249Q) and peptidase-inactive (D493A) variants are also shown. (B) Peptidase activity of recombinant *Pf*FtsH1 WT and mutants. Data are shown as mean ±s.e.m. (n=3). (C) ATPase activity of recombinant *Pf*FtsH1 WT and mutants. Data are shown as mean ± s.e.m. (n=3).

We first measured peptidase activity on a fluorescent casein substrate. Neither the I437S nor D493A variant had detectable peptidase activity, whereas WT enzyme turned over substrate at 0.4 min^-1^ (Figure 3B). As expected, WT and E249Q showed robust ATP-dependent proteolytic activity (Figure 3B). Similarly, the I437S and E249Q variants no detectable ATP hydrolysis activity, in contrast to both WT and the D493A variant (Figure 3C). Taken together, these results validate the identified missense mutation leading to expression of inactive *Pf*FtsH1 I437S variant as causative for apicoplast loss.

### *Pf*Atg7, a member of the Atg8 conjugation pathway, is a cytoplasmic protein required for apicoplast biogenesis

Though non-apicoplast proteins are expected to play a role in apicoplast biogenesis, all apicoplast biogenesis proteins validated so far have localized to the apicoplast since this criteria is most often used to select candidates. A significant advantage of our forward genetic screen is that it can uncover non-apicoplast pathways required for apicoplast biogenesis, which are biased against by other approaches. A cytoplasmic protein strongly identified in our screen was *Pf*Atg7 (Pf3D7_1126100), which contained a nonsense mutation causing a protein truncation at position Q719. The premature stop codon was upstream of the E1-like activating domain, consistent with *Pf*Atg7 loss-of-function (Figure 4A). In yeast and mammalian cells, Atg7 is required for conjugation of autophagy-related protein 8 (Atg8) to the autophagosome membrane (Ichimura et al., 2000). *Pf*Atg7 has not specifically been implicated in apicoplast biogenesis; however, *Pf*Atg8 has been shown to localize to the apicoplast membrane and be required for apicoplast biogenesis. In analogy with its role in autophagy, *Pf*Atg7 is likely required for conjugating *Pf*Atg8 to the apicoplast membrane. Therefore we suspected the loss-of-function mutant we identified caused apicoplast loss via loss of membrane-conjugated *Pf*Atg8 (Walczak et al., 2018).

**Figure 4.**
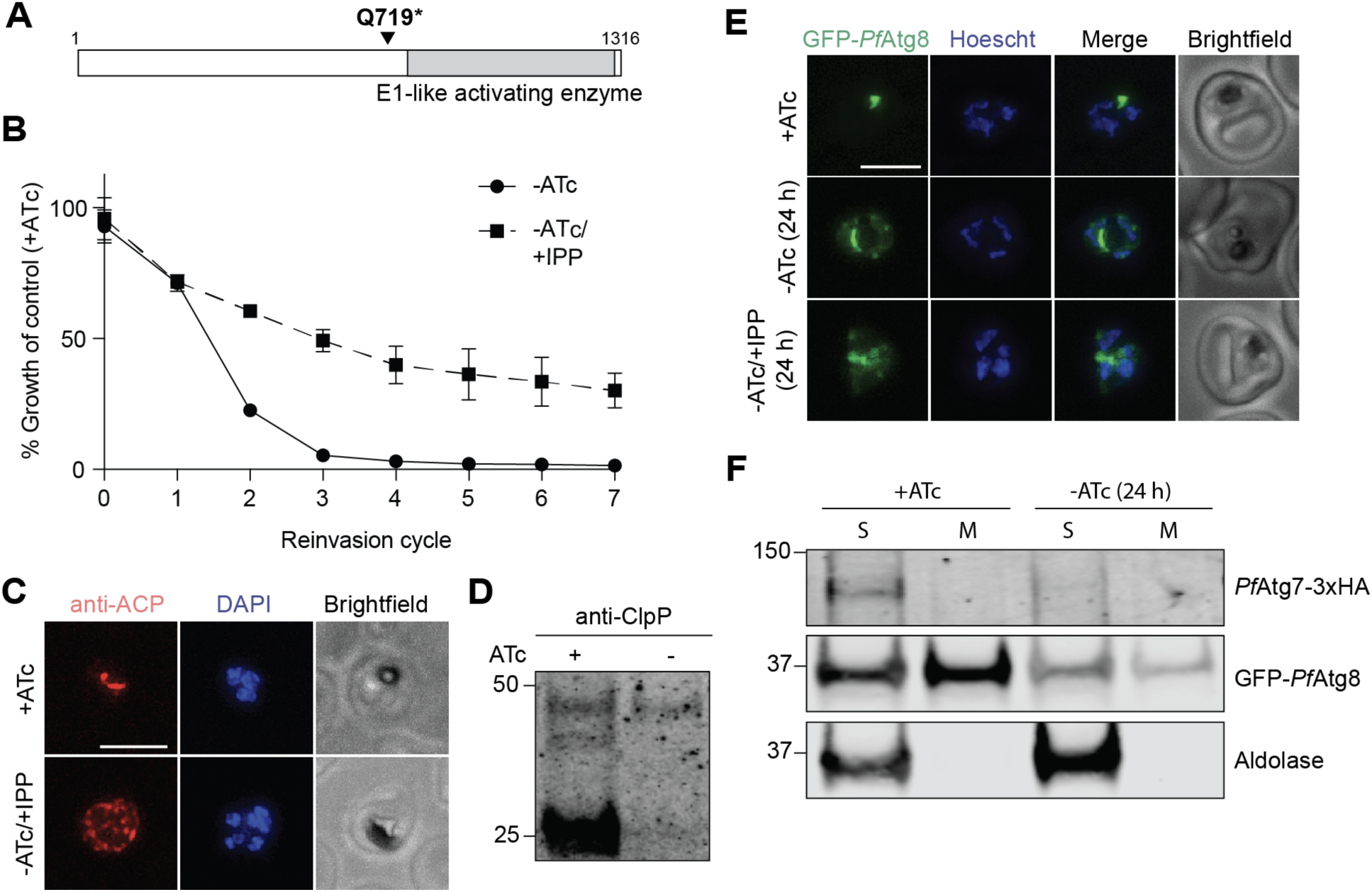
*Pf*Atg7 is a cytoplasmic protein required for *Pf*Atg8 membrane conjugation and apicoplast biogenesis. (A) Domain organization of *Pf*Atg7 showing location of the identified STOP codon. (B) Growth time course of *Pf*Atg7-TetR/DOZI knockdown parasites in the absence of ATc with and without IPP. Data are shown as mean ± s.d. (n=2). (C) Representative immunofluorescence images of *Pf*Atg7-TetR/DOZI parasites showing apicoplast morphology (anti-ACP) in +ATc and -ATc/+IPP parasites at reinvasion cycle 7. Scale bar 5 µm. (D) Western blot of apicoplast-targeted ClpP in *Pf*Atg7-TetR/DOZI parasites in the presence and absence of ATc at reinvasion cycle 7. Processed ClpP after removal of its transit peptide is ∼25 kDa, while full-length protein is ∼50 kDa. (E) Live-cell fluorescent images of GFP-*Pf*Atg8 upon *Pf*Atg7 knockdown 24 h post-ATc removal, prior to apicoplast loss. Scale bar 5 µm. (F) Phase separation (Tx-114) of GFP-*Pf*Atg8 and *Pf*Atg7-3xHA 24 h post-ATc removal, prior to apicoplast loss. Aldolase serves as a soluble protein control.

To confirm *Pf*Atg7’s role in apicoplast biogenesis, we generated a *P. falciparum* strain in which it is conditionally expressed. The endogenous *Pf*Atg7 locus in a NF54 pCRISPR strain was modified with a *C*-terminal triple HA tag and a 3’UTR RNA aptamer sequence that binds a TetR/DOZI repressor to generate *Pf*Atg7-TetR/DOZI (Ganesan et al., 2016; Wagner et al., 2014). In the presence of anhydrotetracycline (ATc), the 3’UTR aptamer is unbound and *Pf*Atg7-3xHA protein was detectable, albeit at low levels consistent with its low expression in published transcriptome data (PlasmoDB.org) (Figure S2A and 4F). Removal of ATc causes binding of the TetR/DOZI repressor, which resulted in undetectable *Pf*Atg7 protein levels by western blot within one replication cycle (Figure S2A and 4F). Parasitemia also decreased over the course of several replication cycles, consistent with a previous study showing that *Pf*Atg7 is essential (Figure 4B) (Walker et al., 2013).

We tested whether the requirement for *Pf*Atg7 was due to its role in apicoplast biogenesis. Growth inhibition caused by *Pf*Atg7 knockdown was partially rescued by addition of IPP (Figure 4B). Furthermore, in –ATc/+IPP parasites, the apicoplast marker acyl-carrier protein (ACP) mislocalized from a discrete organellar localization to multiple cytoplasmic puncta, a hallmark of apicoplast loss (Figure 4C) (Yeh and DeRisi, 2011). Loss of transit peptide cleavage of the apicoplast protein ClpP also confirmed apicoplast loss, since mislocalized apicoplast proteins do not undergo removal of their targeting sequences (Figure 4D) (Gisselberg et al., 2013) (unpublished data). Interestingly, IPP rescue of *Pf*Atg7 knockdown parasites was incomplete, raising the possibility that *Pf*Atg7 is also required for a non-apicoplast function. Alternatively, *Pf*Atg7 knockdown may cause stalling of apicoplast morphologic development leading to general cellular toxicity that cannot be fully rescued with IPP until apicoplast loss is complete. To test these models, instead of *Pf*Atg7 knockdown followed by apicoplast loss, we first induced apicoplast loss with actinonin and then assessed the effects of *Pf*Atg7 knockdown. *Pf*Atg7 knockdown in *apicoplast(-)* parasites did not cause any additional growth defect and was fully rescued by IPP, suggesting that the partial rescue observed in *apicoplast(+)* parasites was due to the order of disruption (Figure S3). These results confirmed that *Pf*Atg7 is required for apicoplast biogenesis and likely is its only essential function.

Finally, to determine whether *Pf*Atg7’s role in apicoplast biogenesis was via *Pf*Atg8 membrane conjugation, we transfected *Pf*Atg7-TetR/DOZI with a transgene encoding GFP-*Pf*Atg8. In this strain, GFP-*Pf*Atg8 primarily localized to a branched structure in schizonts, consistent with its previously described apicoplast localization (Figure 4E) (Kitamura et al., 2012; Tomlins et al., 2013). Upon *Pf*Atg7 knockdown, GFP-*Pf*Atg8 localization to this discrete structure was lost and accumulation in the cytoplasm was observed within a single replication cycle prior to apicoplast loss (Figure 4E). Membrane fractionation also showed an increased proportion of soluble GFP-*Pf*Atg8, while *Pf*Atg7 remained solely in the soluble fraction as expected for a cytoplasmic protein (Figure 4F). These results suggest that, like yeast and mammalian Atg7 homologs, *Pf*Atg7 has a conserved function in conjugating Atg8 to lipids. Altogether, *Pf*Atg7 stood out as a cytoplasmic protein required for apicoplast biogenesis identified in our screen.

### Forward genetics identifies three “conserved *Plasmodium* proteins of unknown function” required for apicoplast biogenesis

The real power of forward genetics is the ability to uncover novel pathways without any *a priori* knowledge. Therefore we next turned our attention to the nearly 50% of the *Plasmodium* genome annotated as “conserved *Plasmodium* protein, unknown function.” Three candidate genes (Pf3D7_0518100, Pf3D7_1305100, Pf3D7_1363700) encoding proteins of unknown function were identified in our screen. Incidentally, all were also identified in a proteomic dataset of apicoplast proteins (unpublished data). Each candidate harbored nonsense mutations that caused protein truncation. The position of the premature stop codon near the 5’ end (Pf3D7_0518100, Pf3D7_1305100) or in the middle (Pf3D7_1363700) of the genes suggested that these were loss-of-function mutations (Figure 5A).

**Figure 5.**
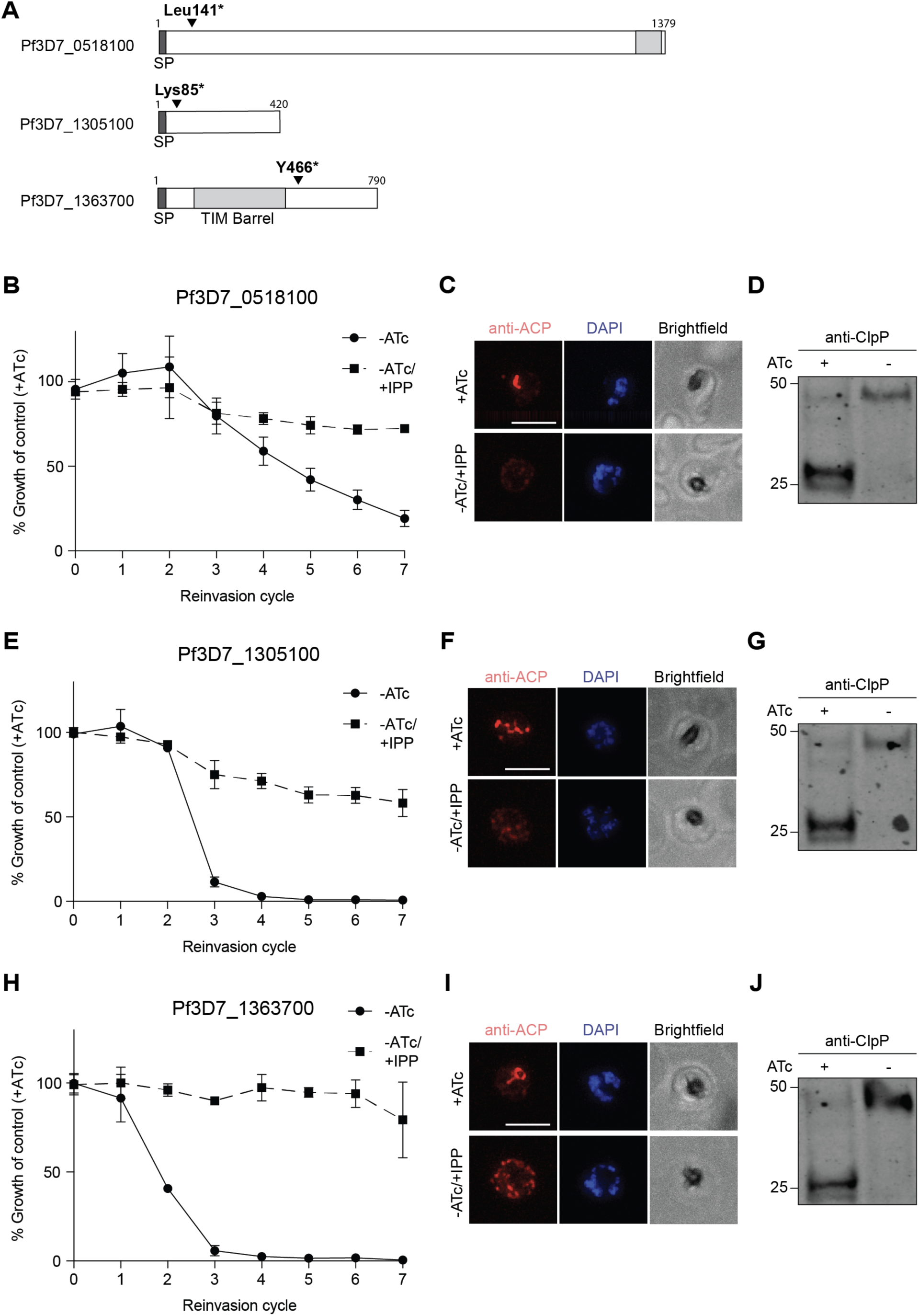
Three “conserved *Plasmodium* proteins of unknown function” are required for apicoplast biogenesis. (A) Domain organization of Pf3D7_0518100, Pf3D7_1305100, and Pf3D7_1363700 showing location of identified STOP codons. (B, E, and H) Growth time course of Pf3D7_0518100, Pf3D7_1305100, and Pf3D7_1363700-TetR/DOZI knockdown parasites in the absence of ATc with and without IPP. Data are shown as mean ± s.d. (n=2). (C, F, and I) Representative immunofluorescence images of indicated TetR/DOZI parasites showing the apicoplast (anti-ACP) in +ATc and -ATc/+IPP parasites at reinvasion cycle 7. Scale bars 5 µm. (D, G, and J) Western blot of apicoplast-targeted ClpP in indicated TetR/DOZI parasites in the presence and absence of ATc at reinvasion cycle 7. Processed ClpP after removal of its transit peptide is ∼25 kDa, while full-length protein is ∼50 kDa.

Therefore we assessed whether knockdown of these genes disrupted apicoplast biogenesis. Similar to the experiments performed to validate *Pf*Atg7, ATc-regulated knockdown strains for each of the candidate genes were generated. Upon ATc removal, protein levels for each gene decreased within 24 hours as detected by western blot (Figure S2B-D). Significant growth inhibition was also observed for all candidate genes tested, with varying kinetics of growth inhibition observed for each candidate (Figure 5B, E, and H). Of note, the gene essentiality demonstrated here for Pf3D7_1305100 and Pf3D7_1363700 confirmed whole-genome essentiality data reported in *P. berghei* and/or *P. falciparum.* However, the essentiality of Pf3D7_0518100 did not agree with its “dispensable” annotation in the *P. falciparum* dataset. IPP supplementation reversed growth inhibition for all the candidates, demonstrating that their essentiality was due to an apicoplast-specific function (Figure 5B, E, and H). Finally, mislocalization of ACP and loss of transit peptide cleavage of ClpP confirmed apicoplast loss for all candidates (Figure 5C-D, F-G, I-J). As these genes have so far lacked any functional annotation and given their shared knockdown phenotype, we designated them “<u>a</u>picoplast-<u>m</u>inus, IPP-<u>r</u>escued” (*amr*) genes: *amr1* (Pf3D7_1363700), *amr2* (Pf3D7_0518100), and *amr3* (Pf3D7_1305100). Taken together, we successfully identified three novel proteins of unknown function required for apicoplast biogenesis, prioritizing these *amr* genes for functional studies.

### Sequence analysis of AMR1 homologs suggests gene duplication followed by evolution of a new function in apicoplast biogenesis

To set up future functional studies, we noted that *Pf*AMR1 contained a TIM-barrel domain with closest homology to indole-3-glycerol phosphate synthase (IGPS), a highly-conserved enzyme in the tryptophan (trp) biosynthesis pathway (Imanian and Keeling, 2014; Jiroutova et al., 2007; Nagano et al., 2002). This was surprising because *Plasmodium* and the related apicomplexan parasite *Toxoplasma* are trp auxotrophs (Liu et al., 2006; Pfefferkorn, 1984; Sibley et al., 1994). Indeed, analysis of >30 apicomplexan genomes did not detect any of the other six trp biosynthetic enzymes, except the terminal enzyme trpB which was horizontally transferred into *Cryptosporidium* spp. (Huang et al., 2004). Therefore we suspected that *Pf*AMR1 may have a function unrelated to trp biosynthesis.

To test the conservation of active site residues, we aligned the sequences of several known IGPSs with IGPS homologs identified from *P. falciparum, T. gondii*, and *V. brassicaformis* (Figure S4) (Jones, 1999; Pei et al., 2008; Slabinski et al., 2007). *V. brassicaformis*, a member of the Chromerids, is the closest free-living, photosynthetic relative to apicomplexan parasites. It contains a secondary plastid with the same origin as the apicoplast, but as a free-living algae (Moore et al., 2008), is also expected to have intact trp biosynthesis. Known catalytic and substrate binding residues based on enzyme structure-function studies performed in bacteria were first identified (Hennig et al., 2002). For known IGPSs and one of the *V. brassicaformis* IGPS homologs (Vbra_4894), the key catalytic and substrate binding residues were all conserved, despite the vast evolutionary distance between bacteria, metazoans, and Chromerids/apicomplexans (Figure 6A). However, in *Pf*AMR1 and the other two *V. brassicaformis* homologs, key functional residues were not conserved. Based on the conservation of functional residues, we separated these sequences into 2 groups: “canonical IGPS” sequences that have been shown or are likely to encode for IGPS activity and “IGPS-like” sequences (e.g. *Pf*AMR1) which we suggest have functionally diverged.

**Figure 6.**
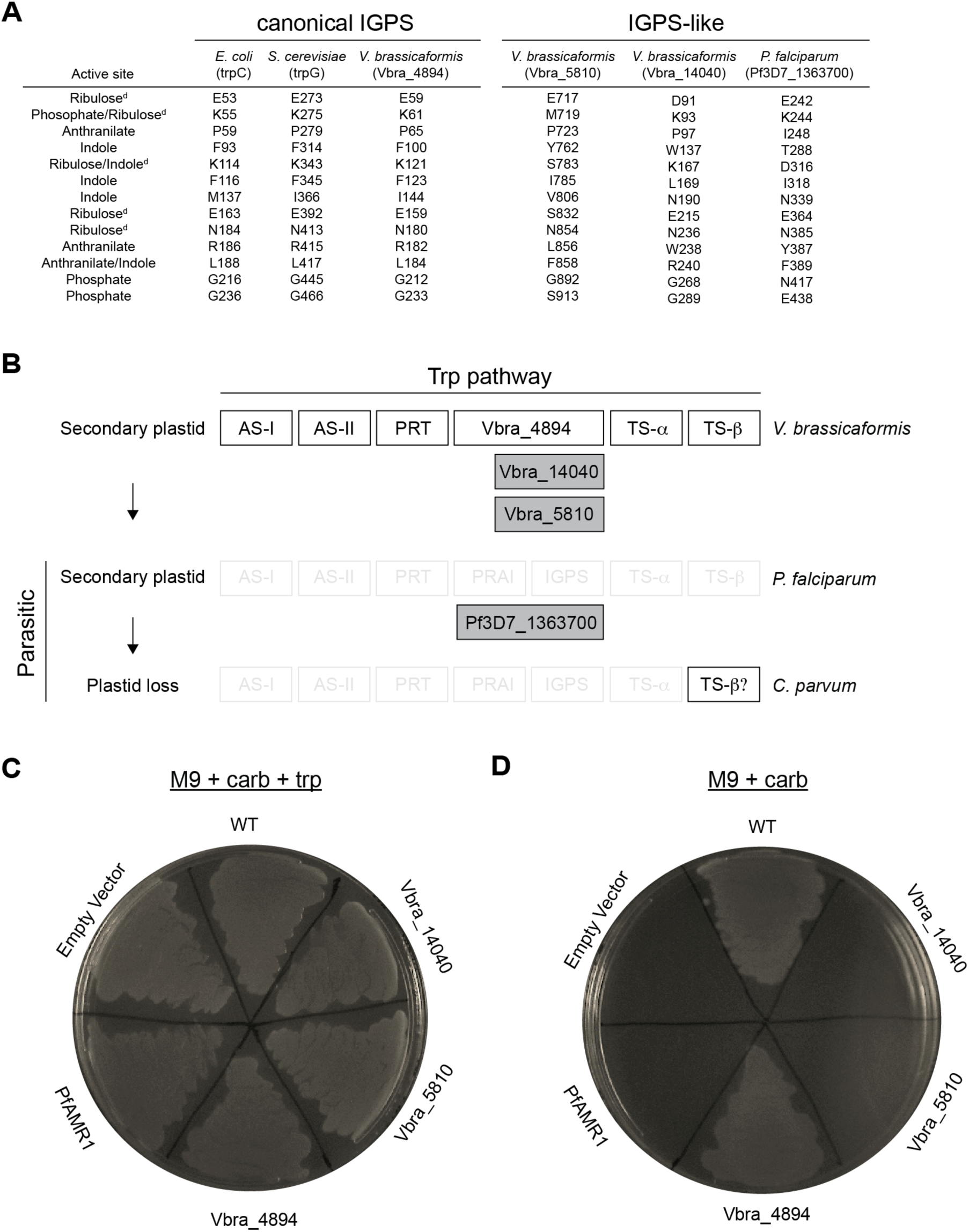
Evolutionary divergence from IGPS proteins suggest a novel apicoplast biogenesis function for *Pf*AMR1. (A) Conservation of of active-site residues between IGPS and IGPS-like proteins. Sequences of IGPS homologs identified in *V. brassicaformis* and *P. falciparum* and known IGPS enzymes from *E. coli* and *S. cerevisiae* were aligned. Active-site residues identified from structure-function studies in IGPS enzymes are listed. ^d^ Catalytic residues. See also Figure 4. (B) Schematic of IGPS and IGPS-like protein loss in *P. falciparum* and *C. parvum*, respectively. (C) Complementation of an *E. coli* trpC mutant (*trpC9800*) with IGPS and IGPS-like proteins from *V. brassicaformis* and *P. falciparum* grown on minimal media (M9) + antibiotic resistance (carbenicillin) + and tryptophan (trp). Empty vector condition contains only antibiotic resistance. *BL21 Star (DE3)* strain was used for the WT condition. (D) Same complementation as in (C) but grown without trp.

We next looked at the pattern of canonical IGPS versus IGPS-like proteins through two biological transitions: loss of trp biosynthesis (*Vitrella* vs *Plasmodium* spp.) and loss of the apicoplast (*Plasmodium* vs *Cryptosporidium* spp.) (Figure 6B). As expected for a role in trp biosynthesis, canonical IGPS proteins were retained until the emergence of parasitism. In addition, genes encoding the remaining set of enzymes for trp biosynthesis were identified in the *V. brassicaformis* genome (Woo et al., 2015). Unlike canonical IGPSs however, IGPS-like proteins were retained in parasites that have lost trp biosynthesis. Instead, loss of IGPS-like proteins is associated only with loss of the apicoplast in *Cryptosporidium* spp. This pattern of acquisition and loss of IGPS-like proteins suggests an apicoplast-specific function separate from trp biosynthesis.

Finally, we performed functional complementation to test the biochemical activity of canonical IGPS and IGPS-like genes from *V. brassicaformis* and *P. falciparum*. An *E. coli* strain (*trpC9800*) containing an inactivating mutation in trpC, the *E. coli* IGPS homolog, was grown on minimal agar (M9) (Yanofsky et al., 1971). As expected, *trpC9800* was dependent on trp supplementation for growth (Figure 6C). Complementation with the Vbra_4894 homolog also restored *trpC9800* growth on M9, comparable to that of an *E. coli* strain with intact tryptophan biosynthesis, suggesting Vbra_4894 is an IGPS protein (Figure 6D). In contrast, neither *Pf*AMR1 nor any of the IGPS-like genes were able to functionally replace trpC, supporting an alternative biochemical function.

## Discussion

Apicoplast biogenesis provides a fascinating window into molecular evolution, including examples of proteins that have been reused (e.g. TIC/TOC complexes) (Kalanon et al., 2009; Sheiner et al., 2015; van Dooren et al., 2008), repurposed (e.g. Atg8, SELMA) (Spork et al., 2009; Walczak et al., 2018), or newly invented in the process of serial endosymbioses. Overcoming significant technical challenges in the *Plasmodium* experimental system, we designed a forward genetic screen to identify essential apicoplast biogenesis pathways. This singular screen opens up opportunities to discover evolutionary innovations obscured by candidate-based approaches, including cytoplasmic pathways and genes lacking any functional annotations. In addition to confirming the role of *Pf*FtsH1 in apicoplast biogenesis and identification of *Pf*Atg8 conjugation machinery, we identified several proteins of unknown function required for apicoplast biogenesis that have so far gone undetected.

One surprising gene we identified was *Pf*AMR1 which encodes a TIM-barrel domain found in diverse enzymes catalyzing small molecule reactions (Nagano et al., 2002). *Pf*AMR1 may have evolved from gene duplication of indole-3-glycerol phosphate synthase (IGPS), an enzyme in the tryptophan biosynthetic pathway (Jiroutova et al., 2007). However, the evolutionary pattern of retention in apicomplexan parasites lacking trytophan biosynthesis and loss in *Cryptosporidium* spp., concomitant with plastid loss, supports a critical function of AMR1 in the apicoplast independent of trytophan biosynthesis. We hypothesize that *Pf*AMR1 may be involved in biosynthesis of a specialized lipid or signaling molecule required specifically for building new plastids in this lineage. Multiple new AMR genes identified in this study provide striking examples of the unexpected findings enabled by unbiased screens.

Uncovering novel apicoplast biogenesis pathways also has important biomedical applications. While targeting the metabolic function of the apicoplast has been a major strategy for antimalarial drug discovery (Jomaa et al., 1999), it has become apparent that apicoplast biogenesis is equally or likely more valuable as a therapeutic target (Amberg-Johnson et al., 2017). These distinct pathogen pathways are nonetheless required in every proliferative stage of the *Plasmodium* lifecycle and conserved among apicomplexan parasites. Targeting apicoplast biogenesis has the benefit of efficacy against multiple *Plasmodium* life stages and multiple pathogens. Consistent with this broad utility, antibiotics that inhibit translation in the apicoplast and disrupt its biogenesis are used clinically for malaria prophylaxis, acute malaria treatment, and treatment of babesiosis and toxoplasmosis (Dahl and Rosenthal, 2007; Fichera and Roos, 1997; Goodman et al., 2016).

Until now, a forward genetic screen for essential pathways in blood-stage *Plasmodium* has not been achieved. Previous screens in murine *P. berghei* and the human malaria parasite *P. falciparum* identified non-essential genes required for gametocyte formation (Ikadai et al., 2013; Sinha et al., 2014), the developmental stage required for mosquito transmission. Functional screens for essential pathways have been impeded by several technical challenges, including the low transfection efficiency of *P. falciparum, in vivo* growth requirement of *P. berghei*, and absence of efficient methods for generating conditional mutants in both organisms (de Koning-Ward et al., 2015). Nonetheless, genome-scale deletion screens in *P. berghei* and *P. falciparum* using a homologous recombination-targeted deletion library or saturating transposon-based mutagenesis, respectively, have revealed a plethora of essential genes (Bushell et al., 2017; Zhang et al., 2018). Functional assignment of these essential genes is a priority. In this context, the apicoplast biogenesis screen presented here is a major milestone towards unbiased functional identification of novel, essential genes.

A top priority for “version 2.0” is to expand the screen to genome scale, maximizing our ability to uncover novel pathways. Apicoplast biogenesis is a complex process encompassing a multitude of functions and is expected to require hundreds of gene products. The identification of 12 candidates genes in the current screen is far from saturating. The most significant bottleneck is the dependence of this screen on chemical mutagenesis. Mapping mutations by whole-genome sequencing limits the number of mutants that can be analyzed and in our study, less than half had a detectable point mutation. Specific mutations identified in candidate genes also need to be validated one-by-one. In this study, four nonsense mutations were validated by conditional knockdown, while a missense mutation in *Pf*FtsH1 was validated using an available activity assay. Although we were able to demonstrate loss-of-function as result of the *Pf*FtsH1 I437S variant, other missense mutations identified in genes of unknown function will be challenging to follow up with available genetic tools. Given these limitations, an alternative mutagenesis method will increase the screen throughput. Options include adaptation of the *piggyBac* transposon developed for the *P. falciparum* deletion screen (Zhang et al., 2018) or development of large-scale targeted mutagenesis. Switching to more genetically-tractable apicomplexan organisms, such as *P. berghei* or *Toxoplasma*, would provide ready options for large-scale targeted gene disruptions (Bushell et al., 2017; Sidik et al., 2016). However, these would have to be performed under conditional regulation since chemical rescue of the apicoplast is not feasible in these organisms. We anticipate that continued advances in genetic methods in apicomplexan organisms will open up opportunities to expand this screen in the future.

## Acknowledgements

We thank Dr. Sean Prigge for anti-ACP antibody, Dr. Walid Houry for the anti-ClpP antibody, and Dr. Vida Ahyong for guidance on data analysis. Ellen Yeh is a Chan Zuckerberg Biohub Investigator. This project has been funded with federal funds from the NIAID, NIGMS, and Director’s Fund, National Institutes of Health, Department of Health and Human Services under Award Numbers 1K08AI097239 (EY), 1DP5OD012119 (EY), AI016892 (RTS), F32GM116241 (SBH), T32GM007276 (KAJ &TRM), GM007365 (YT), and S10OD018220 (Stanford Functional Genomics Facility). Funding was also provided by the Burroughs-Wellcome Fund (EY), Chan Zuckerberg Biohub (EY), and the Stanford Bio-X SIGF William and Lynda Steere Fellowship (KAJ). The content is solely the responsibility of the authors and does not necessarily represent the official views of the National Institutes of Health.

## Author Contributions

Conceptualization, EY, KAJ, YT; Methodology, YT, TRM, KAJ, MW, MJPG, SBH, RTS, EY; Software, YT and TRM; Formal Analysis, YT, TRM, SBH, MW; Investigation, YT, TRM, MW, MJPG, SBH; Writing – Original Draft, EY and YT; Writing – Review & Editing, EY, YT, SBH, TRM, MW, MJPG, KAJ, RTS; Visualization, YT, MJPG, SBH; Supervision, EY and RTS; Funding Acquisition, EY and RTS.

The authors declare no competing interests.

## Methods

### Parasite culture and transfections

*P. falciparum* Dd2^attB^ (MRA-843) were obtained from MR4. NF54^Cas9+T7 Polymerase^ parasites were kindly provided by Jacquin Niles. Parasites were grown in human erythrocytes at 2% hematocrit (Stanford Blood Center) in RPMI 1640 media (Gibco) and supplemented with 0.25% Albumax II (Gibco), 2 g/L sodium bicarbonate, 0.1 mM hypoxanthine (Sigma), 25 mM HEPES, pH 7.4 (Sigma), and 50 µg/L gentamicin (Gold Biotechnology) at 37°C, 5% O_2_, and 5% CO_2_.

For transfections, 50 µg of plasmid DNA was added to 200 µL packed red blood cells, adjusted to 50% hematocrit in RPMI 1640, and electroporated as previously described (Deitsch et al., 2001). On day 4 post-transfection, parasites were selected for with 2.5. mg/L blasticidin S (RPI Research Products). TetR/DOZI strains were cultured with 500 nM anhydrous tetracycline (ATc) for the entire duration of transfection. For TetR/DOZI strains expressing ACP_L_-GFP or GFP-*Pf*Atg8, parasites were additionally selected for with 500 µg/mL G418 sulfate (Corning) during transfection.

### Cloning

Oligonucleotides were purchased from the Stanford Protein and Nucleic Acid facility or IDT. gBlocks were ordered from IDT. Molecular cloning was performed using In-Fusion cloning (Clontech) or Gibson Assembly (NEB). Primer and gBlock sequences for all cloning are available in Table S3.

To generate the plasmid pRL2-mCherry-T2A-ACP_L_-GFP, T2A-ACP_L_-GFP was first amplified from the pRL2-ACP_L_-GFP vector. mCherry was amplified from pTKO2-mCherry vector (kind gift from J. Boothroyd) and inserted in front of T2A-ACP_L_-GFP in the pRL2 backbone using the In-Fusion Cloning kit (Takara). To generate the pL2-mCherry-T2A-ACP_L_-GFP-degron plasmids, T2A-ACP_L_-GFP-degron was amplified from pRL2-mCherry-T2A-ACP_L_-GFP.

For CRISPR-Cas9-based editing of endogenous Pf3D7_0518100, Pf3D7_1126100, Pf3D7_1305100 and Pf3D7_1363700 loci, sgRNAs were designed using the eukaryotic CRISPR guide RNA/DNA design tool (http://chopchop.cbu.uib.no/). To generate a linear plasmid for CRISPR-Cas9-based editing, left and right homology regions were first amplified for each gene. For each gene, a gBlock containing the recoded coding sequence *C*-terminal of the CRISPR cut site and a triple HA tag was synthesized with appropriate overhangs for Gibson Assembly. This fragment along with the left homology region was simultaneously cloned into the FseI/ApaI sites of the linear plasmid pSN054-V5. Next, the appropriate right homology region and a gBlock containing the sgRNA expression cassette were simultaneously cloned into the AscI/I-SceI sites of the resultant vector to generate the plasmids.

To generate plasmid for expression of GFP-*Pf*Atg8, GFP with a GlyAlaGlyAla linker was amplified from pRL2-ACP_L_-GFP. *Pf*Atg8 was amplified from *P. falciparum* gDNA. Both fragments were inserted into pfYC110 vector (Wagner et al., 2013) using the In-Fusion Cloning kit.

*Vitrella brassicaformis* RNA from strain CCMP3346 was purchased from the National Center for Marine Algae and Microbiota and subsquently used to generate cDNA using Superscript III cDNA Kit (Life Technologies). For *Plasmodium* PF3D7_1363700 cloning, codon optimized gBlocks were used to construct the *Plasmodium* construct. Constructs were cloned into the pGEXT vector using the In-Fusion Cloning kit.

### Degron screening

Ring-stage mCherry-T2A-ACPL-GFP-degron parasites were treated with 10 µM actinonin (Sigma) and 200 µM IPP (Isoprenoids, LLC) to disrupt the apicoplast. Both treated and non-treated parasites were analyzed two cycles post-treatment at the schizont stage on a BD Accuri C6 flow cytometer. 100-500k events were recorded for each condition. Uninfected RBC were first gated away from the infected population. The average fluorescent levels of FL-1A (GFP) and FL-4A (mCherry) were then calculated from each gated population using FlowJo.

### Mutant screening

Ring-stage mCherry-T2A-ACPL-GFP-*Ec*ssrA (*Ec*ssrA) parasites were seeded onto a 96-well plate at a volume of 200 µL, 2% hematocrit and 0.5-1% parasitemia. Parasites were either untreated, treated with 1 mM ethyl methanesulfonate (EMS) or 100 µM N-ethyl-N-nitrosourea (ENU) for 2 h and then washed three times after to remove the mutagen from the growth media. Parasites were cultured in growth media + 200 µM IPP for the duration of the screen.

At 120 h post-treatment, mutants were isolated on a Sony SH800S Cell Sorter. Uninfected RBCs were first analyzed to set the gate for overall fluorescence. Untreated *Ec*ssrA parasites were analyzed to gate for positive and negative mCherry and GFP expression, respectively. Actinonin-treated *Ec*ssrA parasites were analyzed to gate for positive GFP expression. Non-mutagenized and mutagenized parasites displaying both positive mCherry and GFP expression were FACS’d into a new 96-well plate. Enriched parasites were allowed to propagate to a detectable parasitemia before being subjected to subsequent FACS rounds.

Mutants were enriched until mCherry and GFP fluorescence approached actinonin-treated levels. In the final enrichment, up to 52 mutants were single-cell cloned. Mutants that survived single-cell sorting were split into growth media containing either 200 µM IPP or no IPP. Mutants displaying growth only in IPP were expanded to a 10 mL culture at ∼10% parasitemia and ring-stage synchronized. 5 mL of culture was harvested for DNA extraction and 5 mL culture was frozen at −80°C.

### DNA isolation and whole-genome sequencing

Ring-stage parasites were isolated from RBC in 0.1% saponin and washed three times in PBS. gDNA from mutants and the parental strain was isolated using the Quick-DNA Universal Kit (Zymo Research). Paired-end gDNA libraries were generated and barcoded for each mutant and the parental using the Nextera Library Prep Kit, modified for 8 PCR cycles (Stanford Functional Genomics Facility). Up to 26 pooled libraries were analyzed per lane of an Illumina HiSeq 4000 using 2 x 75 bp, paired-end sequencing (Stanford Functional Genomics Facility).

### SNP analysis

Fastq files were checked for overall quality using FastQC. 10 and 15 bp were trimmed from the 5’ and 3’ ends of all 75 bp sequence reads, respectively, to remove low quality reads. 20 and 30 bp were trimmed 5’ and 3’ ends of 150 bp sequence reads, respectively, from an additional parental Dd2 strain sequenced by the DeRisi lab (https://www.ncbi.nlm.nih.gov/sra/SRX326518). The resulting paired-end sequencing reads were mapped using Bowtie2 against the *P. falciparum* 3D7 (version 35) reference genome. One mismatch per read was allowed and only unique reads were aligned (reads were removed if they aligned to more than one region of the genome). PCR duplicates were removed using Samtools rmdup and raw SNV’s were called using Samtools mpileup. Indels were not analyzed.

Bcftools was used to generate the raw variant list for parental (allele frequency ≥ 0.05, depth ≥ 1) and mutant (allele frequency ≥ 0.9, depth ≥ 5) strains. Variants found in the parental strain were excluded from the mutant variant list. Variants were filtered to only include protein-coding mutations (missense and nonsense). Mutations found in hypervariable gene families were excluded. Remaining variants that were previously annotated in PlasmoDB were excluded to generate the final variant list. The reported variants were confirmed to meet the filtering requirements using Samtools tview. Mutants containing nonsense mutations were Sanger sequenced to confirm the presence of the mutations prior to genetic validation. The custom script and parameters used for analysis are available at (https://github.com/yehlabstanford/biogenesis_screen).

### Western blotting

Parasites were separated from RBCs by lysis in 0.1% saponin and were washed in PBS. Parasite pellets were resuspended in PBS containing 2X NuPAGE LDS sample buffer and boiled at 95°C for 10 minutes before separation on NuPAGE gels. Gels were transferred onto nitrocellulose membranes using the Trans-Blot Turbo Transfer System (Bio-Rad). Membranes were blocked in 0.1% Hammarsten casein (Affymetrix) in 0.2X PBS with 0.01% sodium azide. Antibody incubations were performed in a 1:1 mixture of blocking buffer and TBST (Tris-buffered saline with Tween-20; 10 mM Tris, pH 8.0, 150 mM NaCl, 0.25 mM EDTA, 0.05% Tween 20). Blots were incubated with primary antibody overnight at 4°C at the following dilutions: 1:20,000 mouse-α-GFP JL-8 (Clontech 632381); 1:20,000 rabbit-α-*Plasmodium* aldolase (Abcam ab207494); 1:1000 mouse-α-HA 2-2.2.14 (Thermo Fisher 26183); 1:1000 guinea pig-α-ATG8 (Josman LLC); 1:1000 rabbit-α-ClpP (kind gift from W. Houry). Blots were washed three times in TBST and were incubated for 1 hour at room temperature in a 1:10,000 dilution of the appropriate fluorescent secondary antibodies (LI-COR Biosciences). Blots were washed twice in TBST and once in PBS before imaging on a LI-COR Odyssey imager.

### Fractionation assay

Parasites expressing GFP-*Pf*Atg8 were grown in the presence or absence of ATc for 24 hours. 10 ml cultures were lysed with 0.1 % saponin and washed 3 times with PBS. Parasite pellets were resuspended in ice-cold lysis buffer (1X PBS, 1 % Triton X-114 (Thermo Scientific 28332), 2 mM EDTA, 1x protease inhibitors (Pierce A32955)) and incubated on ice for 30 min. Cell debris were removed by 10 min centrifugation at 16,000 x g, 4°C. Supernatant was transferred to a fresh Eppendorf tube, incubated 2 min at 37 °C to allow phase separation and centrifuged 5 min at 16,000 x g, room temperature. The top (aqueous) layer was transferred to another tube. The interphase was removed to avoid cross-contamination between the layers. The bottom (detergent) layer was resuspended in 1X PBS, 0.2 mM EDTA to equalize the volumes of the two fractions. Both fractions were subjected to methanol-chloroform precipitation, resuspended in PBS containing 2X NuPAGE LDS sample buffer, boiled for 5 min at 95°C and analyzed by western blot as described above.

### Microscopy

For live imaging, parasites were settled onto glass-bottom microwell dishes Lab-Tek II chambered coverglass (Thermo Fisher 155409) in PBS containing 0.4% glucose and 2 µg/mL Hoechst 33342 stain (Thermo Fisher H3570). Cells were imaged with a 100X, 1.4 NA objective on an Olympus IX70 microscope with a DeltaVision system (Applied Precision) controlled with SoftWorx version 4.1.0 and equipped with a CoolSnap-HQ CCD camera (Photometrics). Images were captured as a series of z-stacks separated by 0.2 µm intervals, deconvolved (except for mCherry images), and displayed as maximum intensity projections. Brightness and contrast were adjusted equally in SoftWorx or Fiji (ImageJ) for display purposes.

For immunofluorescence, fixed-cell imaging, parasites were first fixed with 4% paraformaldehyde (Electron Microscopy Science 15710) and 0.0075% glutaraldehyde in PBS (Electron Microscopy Sciences 16019) for 20 minutes. Cells were washed once in PBS and allowed to settle onto poly-L-lysine-coated coverslips (Corning) for 60 min. Coverslips were then washed once with PBS, permeabilized in 0.1% Triton X-100/PBS for 10 min, and washed twice more in PBS. Cells were treated with 0.1 mg/mL NaBH_4_/PBS for 10 min, washed once in PBS, and blocked in 5% BSA/PBS. Primary antibodies were diluted in 5% BSA/PBS at the following concentrations: 1:500 rabbit-α-*Pf*ACP (kind gift from S. Prigge), 1:100 rat-α-HA 3F10 (Sigma 11867423001). Coverslips were washed three times in PBS, incubated with secondary antibodies goat-α-rat 488 (Thermo Fisher A-11006) and donkey-α-rabbit (Thermo Fisher A10042) at 1:3000 dilution, and washed three times in PBS prior to mounting in ProLong Gold antifade reagent with DAPI (Thermo Fisher).

### Knockdown assays

Ring-stage TetR/DOZI strain parasites were washed two times in growth media to remove ATc. Parasites were divided into three cultures supplemented with 500 nM ATc, no ATc, or no ATc + 200 µM IPP. Samples were collected at the schizont stage in each growth cycle for flow cytometry analysis and western blot. Parasites in each condition were diluted equally every growth cycle for up to six growth cycles.

For parasitemia measurements, parasite-infected or uninfected RBCs were incubated with the live-cell DNA stain dihydroethidium (Thermo Fisher D23107) for 30 min at a dilution of 1:300 (5 mM stock solution). Parasites were analyzed on a BD Accuri C6 flow cytometer and 100k events were recorded.

### Protein expression and purification

The parent His6-SUMO-*Pf*FtsH191-612-GST plasmid as well as E249Q and D493A mutants were obtained from laboratory stocks. An I437S mutant was constructed by site-directed mutagenesis. Recombinant proteins were expressed from these plasmids and purified as described (Amberg-Johnson et al., 2017).

### Enzyme assays

Rates of ATP hydrolysis by *Pf*FtsH1 were measured using a coupled spectrophotometric assay (Norby, 1988) in protein degradation (PD) buffer (25 mM HEPES, pH 7.5, 200 mM NaCl, 5 mM MgSO4, 10 µM ZnSO4, 10% glycerol) with 3% dimethyl sulfoxide (DMSO) or 50 µM actinonin in 3% DMSO at 37 °C. Protein degradation rates were measured by incubating *Pf*FtsH1 (1 µM) with FITC-labeled (2 µM, Sigma C0528) and unlabeled casein (8 µM) in PD buffer plus 3% DMSO or 50 µM actinonin in 3% DMSO. Reactions were started by adding ATP (4 mM) or buffer with a regeneration system (16 mM creatine phosphate and 75 µg/mL creatine kinase), and degradation was followed by measuring the fluorescence intensity (excitation 485 nm; emission 528 nm) at 37°C.

### Alignment of IGPS and IGPS-like proteins

IGPS and IGPS-like proteins from *V. brassicaformis* were identified by BLAST through using CryptoDB. First, secondary structure was predicted using PSI-PRED in the XtalPred suite (Jones, 1999; Slabinski et al., 2007). Only the sequences containing the TIM barrels of each sequence was used for alignment as there are large N- and C-terminal extensions in the non-canonical proteins. PROMALS3D was subsequently used to perform a multiple sequence alignment based on secondary structure and homology to proteins with determined 3D structures (Pei et al., 2008).

### Bacterial complementation assay

*W3110trpC9800 E. coli* strain was purchased from the Yale University Coli Genetic Stock Center and were made chemically competent using calcium chloride. *BL21 Star (DE3)* competent cells (Thermo Fisher) were used for the WT condition. The competent *W3110trpC9800* cells were transformed with the pGEXT vectors containing the different *Vitrella* or *Plasmodium* IGPS and IGPS-like genes and plated on LB agar plates containing carbenicillin. For each construct, a colony was picked and washed in M9 minimal media (M9) [22 mM potassium phosphate monobasic, 22 mM sodium phosphate dibasic, 85 mM sodium chloride, 18.7 mM ammonium chloride, 2 mM magnesium sulfate, 0.1 mM calcium chloride, and 0.4% glycerol], resuspended in M9, and streaked onto M9/agar plates containing either carbenicillin (100 µg/mL) or carbenicillin and 1mM L-tryptophan (Sigma). Plates were incubated at 37°C and allowed to grow for two days, after which images of the plates were taken.

## Supplemental Information

**Figure S1.**
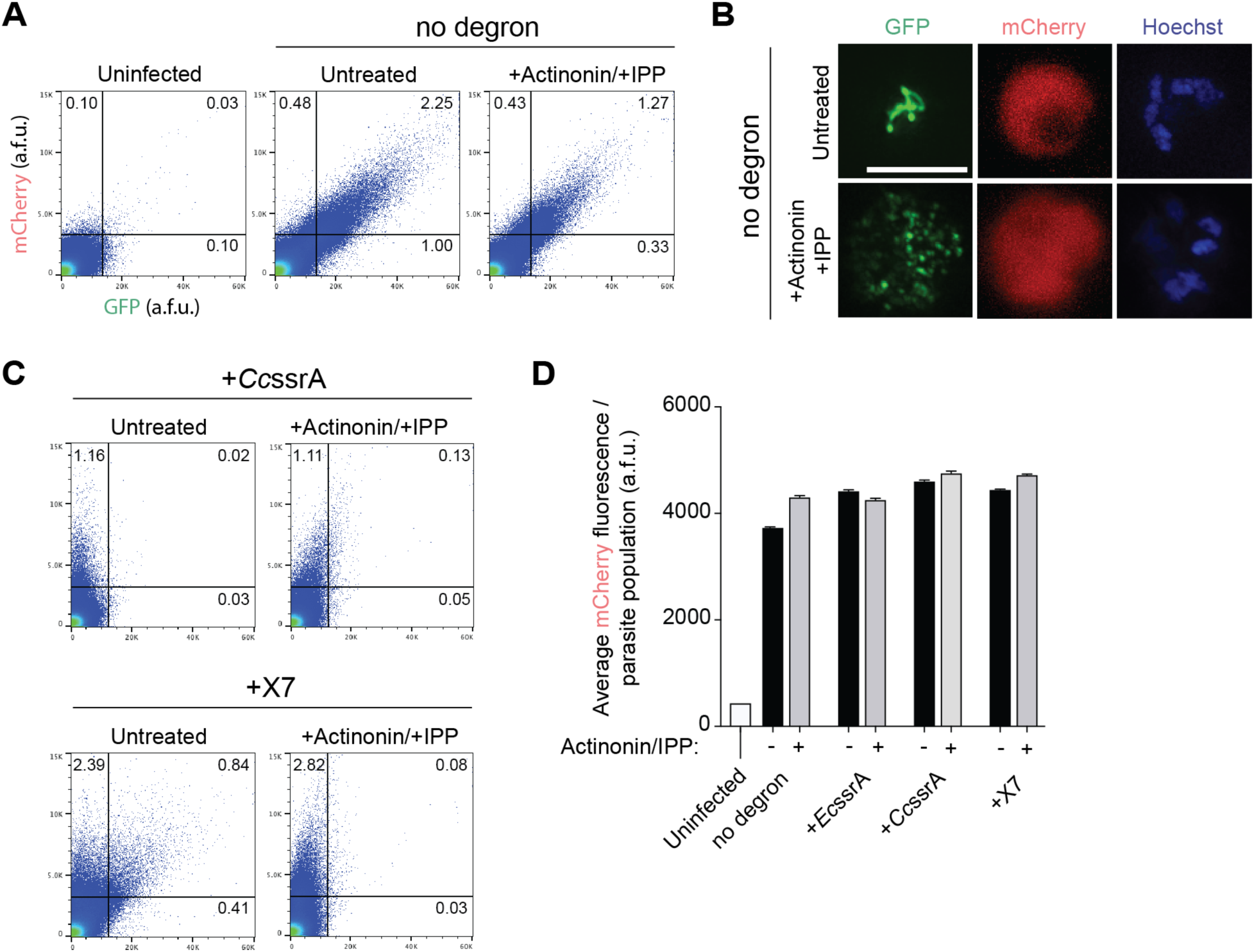
Fluorescence assays for apicoplast reporters. Related to Figure 1. (A) Flow cytometry plots showing mCherry and GFP fluorescence in untreated versus actinonin/IPP treated parasites expressing ACP_L_-GFP with no degron tag. The % mCherry+, GFP+ parasites in the population is indicated. Uninfected red blood cells were used to set gates for mCherry and GFP fluorescence. (B) Representative live-cell fluorescent images of untreated and actinonin/IPP treated parasites expressing mCherry and ACP_L_-GFP. Hoechst stains for parasite nuclei. Scale bar 5 µm. (C) Flow cytometry plots showing mCherry and GFP fluorescence in untreated versus actinonin/IPP treated parasites expressing ACP_L_-GFP-*Cc*ssrA or ACP_L_-GFP-X7. The % mCherry+, GFP+ parasites in the population is indicated. Uninfected red blood cells were used to set gates. (D) Average mCherry fluorescence of reporter strain populations. Data are shown as mean ± s.e.m. (n=2).

**Figure S2.**
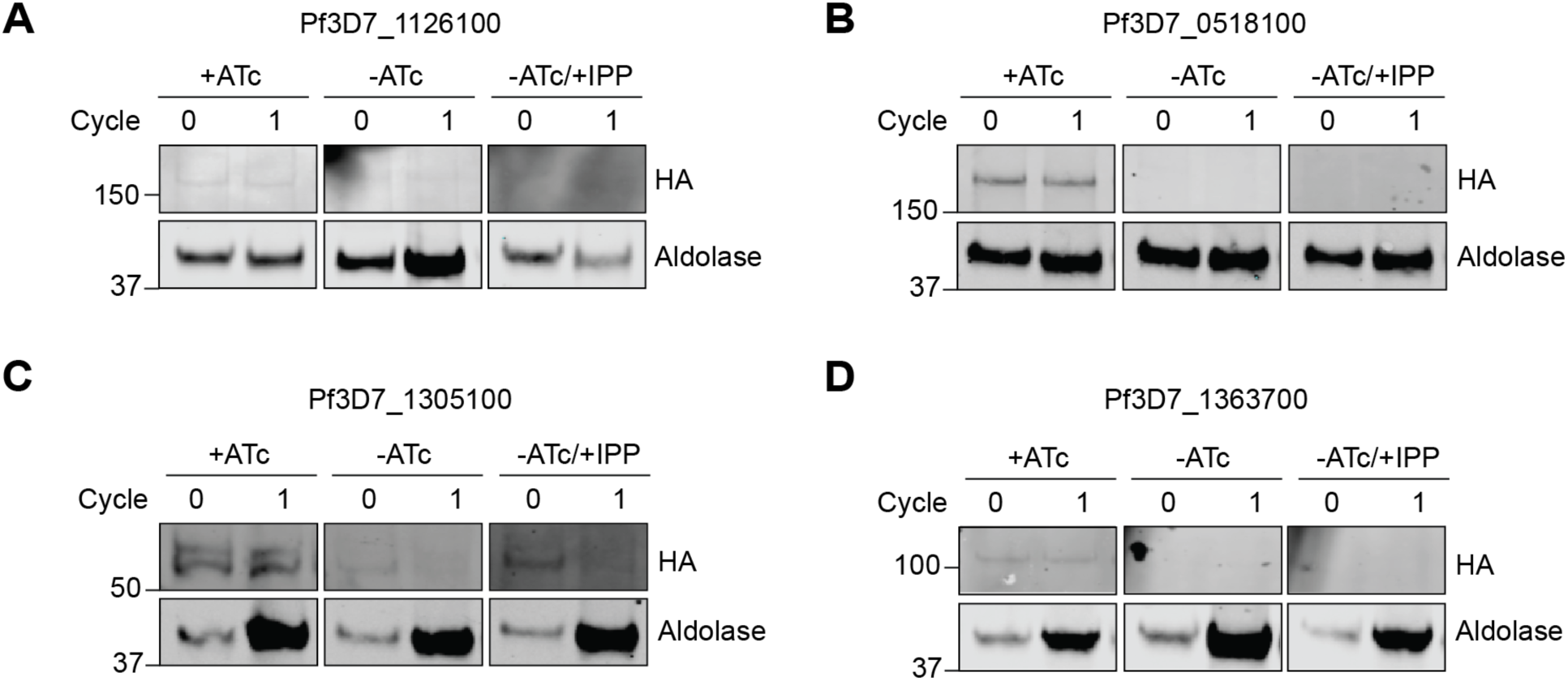
ATc-dependent protein levels in candidate TetR/DOZI parasite strains. Related to Figure 4 and Figure 5. Western blot of HA-tagged protein candidates in TetR/DOZI parasite strains in +ATc, -ATc and -ATc/+IPP parasites. Protein levels for the initial and first reinvasion cycles are shown (0 and 1, respectively). Aldolase serves as a loading control. (A) Pf3D7_1126100 (Atg7), (B) Pf3D7_0518100 (conserved unknown), (C) Pf3D7_1305100 (conserved unknown) and (D) Pf3D7_1363700 (conserved unknown).

**Figure S3.**
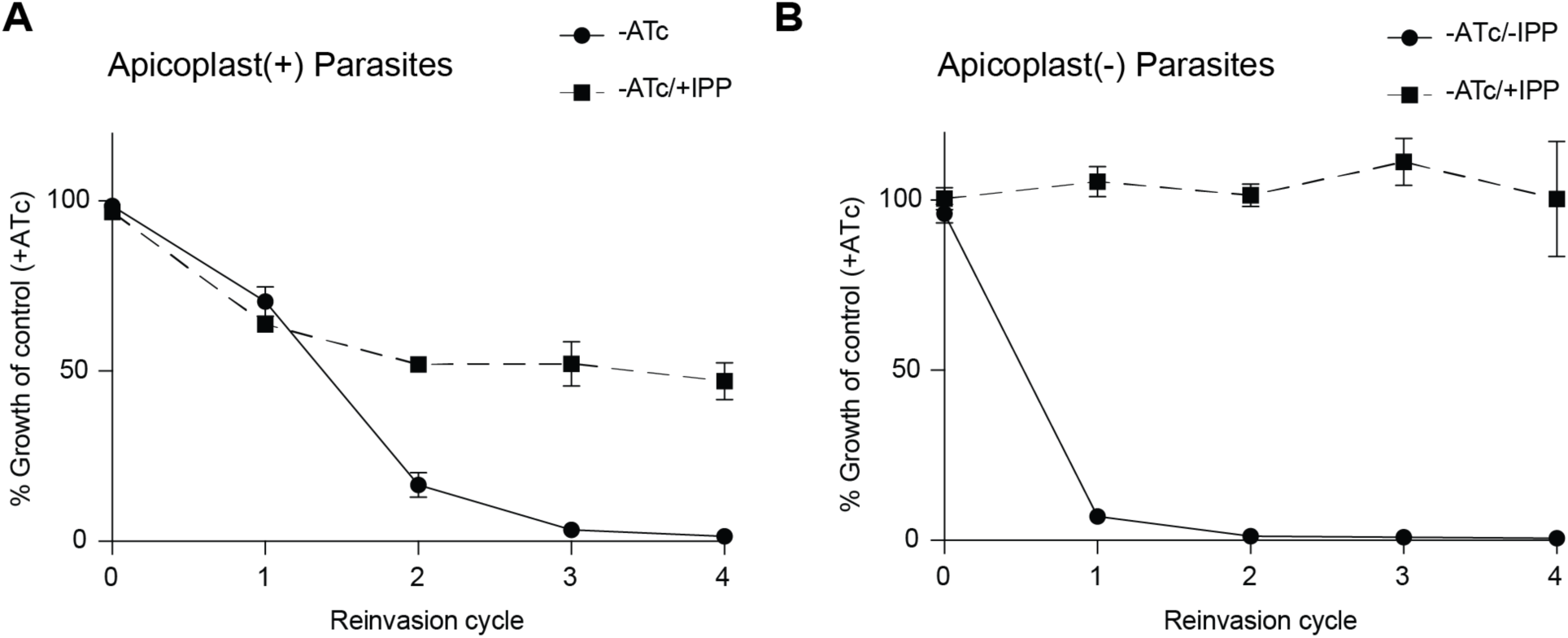
The order of *Pf*Atg7 knockdown and apicoplast loss affects IPP rescue of growth inhibition. Related to Figure 4. Growth time course of *Pf*Atg7-TetR/DOZI parasites in absence of ATc with and without IPP. (A) *Pf*Atg7 knockdown and IPP rescue precedes apicoplast loss in *apicoplast(+)* parasites. Data are shown as mean ± s.d. (n=2). (B) Apicoplast loss precedes *Pf*Atg7 knockdown and IPP in *apicoplast(-)* parasites. *Apicoplast(-)* parasites were generated via actinonin/IPP treatment. Data are shown as mean ± s.d. (n=2).

**Figure S4.**
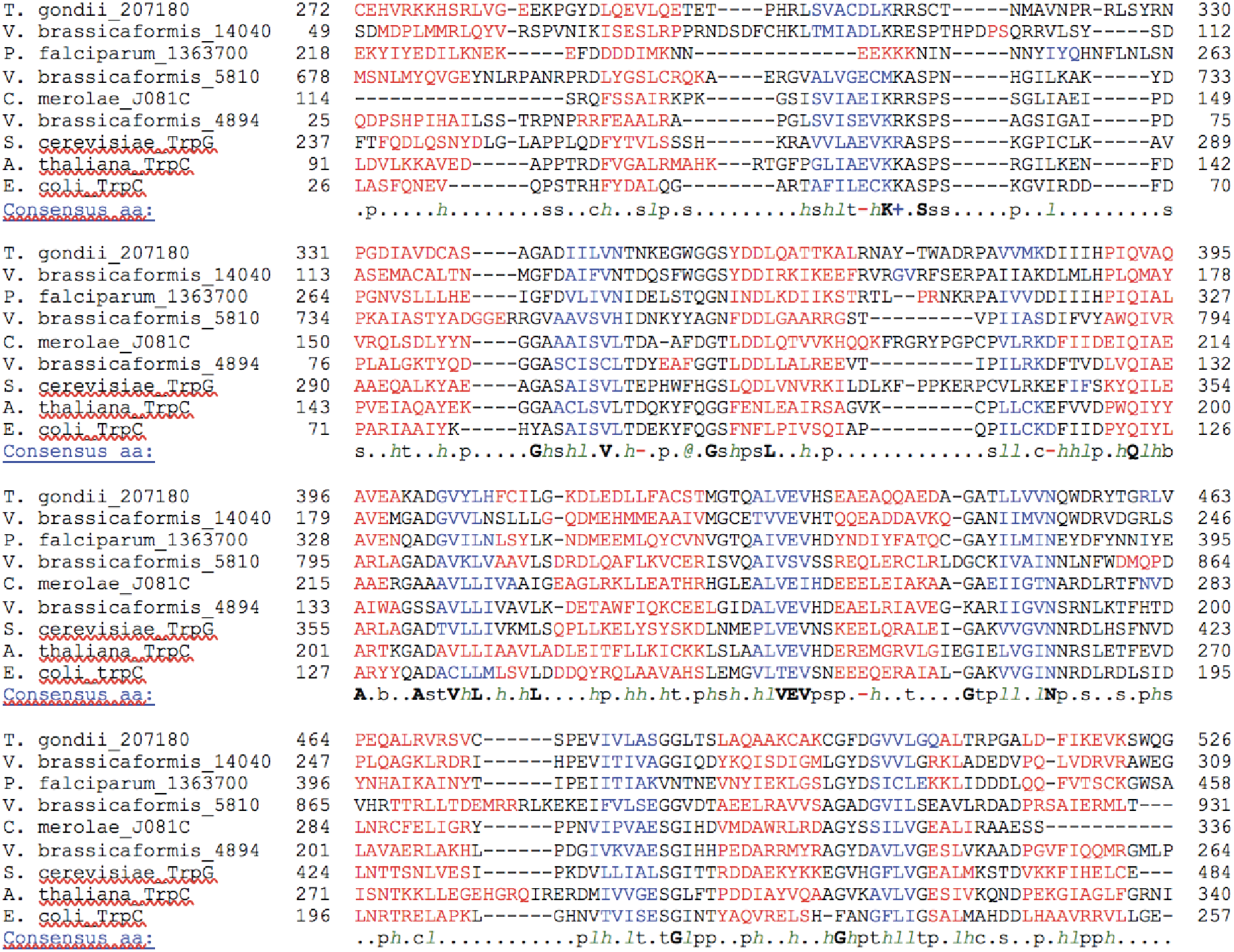
Full sequence alignment of IGPS and IGPS-like proteins from various organisms. Related to Figure 6.

**Table S1.**
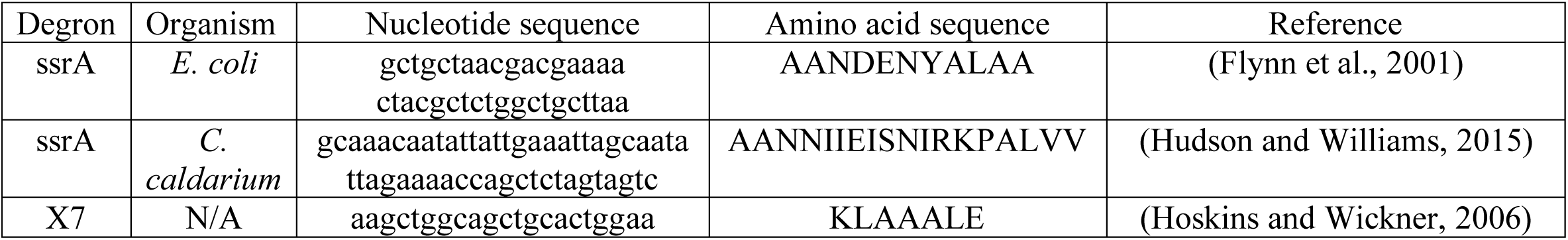
**Amino acid sequences of degrons used for ACP_L_-GFP reporter. Related to Figure 1**

**Table S2. Raw nucleotide variants identified in sequenced clones. Related to Figure 2**

**Table S3. Primers used in this study. Related to Figures 1, 4-6, and S1**

